# Cooperative protein allosteric transition mediated by a fluctuating transmission network

**DOI:** 10.1101/2021.11.17.468836

**Authors:** Matthias Post, Benjamin Lickert, Georg Diez, Steffen Wolf, Gerhard Stock

## Abstract

Allosteric communication between distant protein sites represents a key mechanism of biomolecular regulation and signal transduction. Compared to other processes such as protein folding, however, the dynamical evolution of allosteric transitions is still not well understood. As example of allosteric coupling between distant protein regions, we consider the global open-closed motion of the two domains of T4 lysozyme, which is triggered by local motion in the hinge region. Combining extensive molecular dynamics simulations with a correlation analysis of interresidue contacts, we identify a network of interresidue distances that move in a concerted manner. The cooperative process originates from a cogwheel-like motion of the hydrophobic core in the hinge region, which constitutes a flexible transmission network. Through rigid contacts and the protein backbone, the small local changes of the hydrophobic core are passed on to the distant terminal domains and lead to the emergence of a rare global conformational transition. As in an Ising-type model, the cooperativity of the allosteric transition can be explained via the interaction of local fluctuations.

## INTRODUCTION

Proteins may coexist in various conformational states of different function, e.g., in an active and an inactive state. Transitions between these states can occur spontaneously or may be triggered by an external stimulus such as ligand binding. A prime example is allosteric communication, where a binding event at one site of a protein may affect the structure and dynamics of another distant site.^1–10^ Even without the presence of a ligand, many allosteric systems thermally populate both protein states, thus representing a bistable system. Common to all cases is that the associated structural rearrangement requires the interaction of distant regions of the system. Although these long-range or allosteric couplings appear to be ubiquitous in protein systems,^1^ surprisingly little is known about the dynamical mechanism, its microscopical details, and the time evolution of the associated conformational transition.^8^ We note that the situation is quite different for the protein folding problem, where several decades of theoretical and experimental work have resulted in a quite well-established picture how folding and unfolding proceed. This includes general scenarios such as cooperative two-state and multistate downhill folding,^11^ as well as dynamical mechanisms such as zipping or diffusion limited processes.^12^ The dynamical process underlying an allosteric transition, on the other hand, appears much more elusive due to the typically small local structural changes, which are quite challenging to observe in experiments.^13–15^ While allosteric transitions have been rarely witnessed in unbiased molecular dynamics (MD) simulations,^9,16–20^ numerous residue-based network models have been suggested, which aim to predict intramolecular signaling.^7,21^ Furthermore Markov state models,^22^ which describe the conformational dynamics via memory-less jumps between metastable states, have been applied to allostery.^17,23^ In this work we consider T4 lysozyme (T4L) as a well-studied example of a bistable two-domain protein.^24–29^ Aiming to destroy bacterial cell walls by catalyzing the cleavage of glycosidic bonds, T4L performs an open-closed transition of its two domains that resembles a PacMan, see Fig. 1a. This motion of the “mouth” region was recently shown to be triggered by a locking mechanism in the “hinge” region of T4L, revealing that these two distant regions are allosterically coupled.^29^ (Here we use the term ‘allostery’ for general long-range communication, independent from ligand binding.) With lifetimes of the open and closed state in the order of a few microseconds, T4L is one of the few protein systems, whose large-scale conformational transitions can be sufficiently sampled by unbiased simulations. Adopting the 61 µs-long all-atom MD simulation of Ernst et al.,^29^ here we elucidate the microscopic mechanism of this long-range coupling. Previous work showed that the identification of the underlying reaction coordinates poses a challenge to various dimensionality reduction approaches.^28–31^

**FIG. 1.**
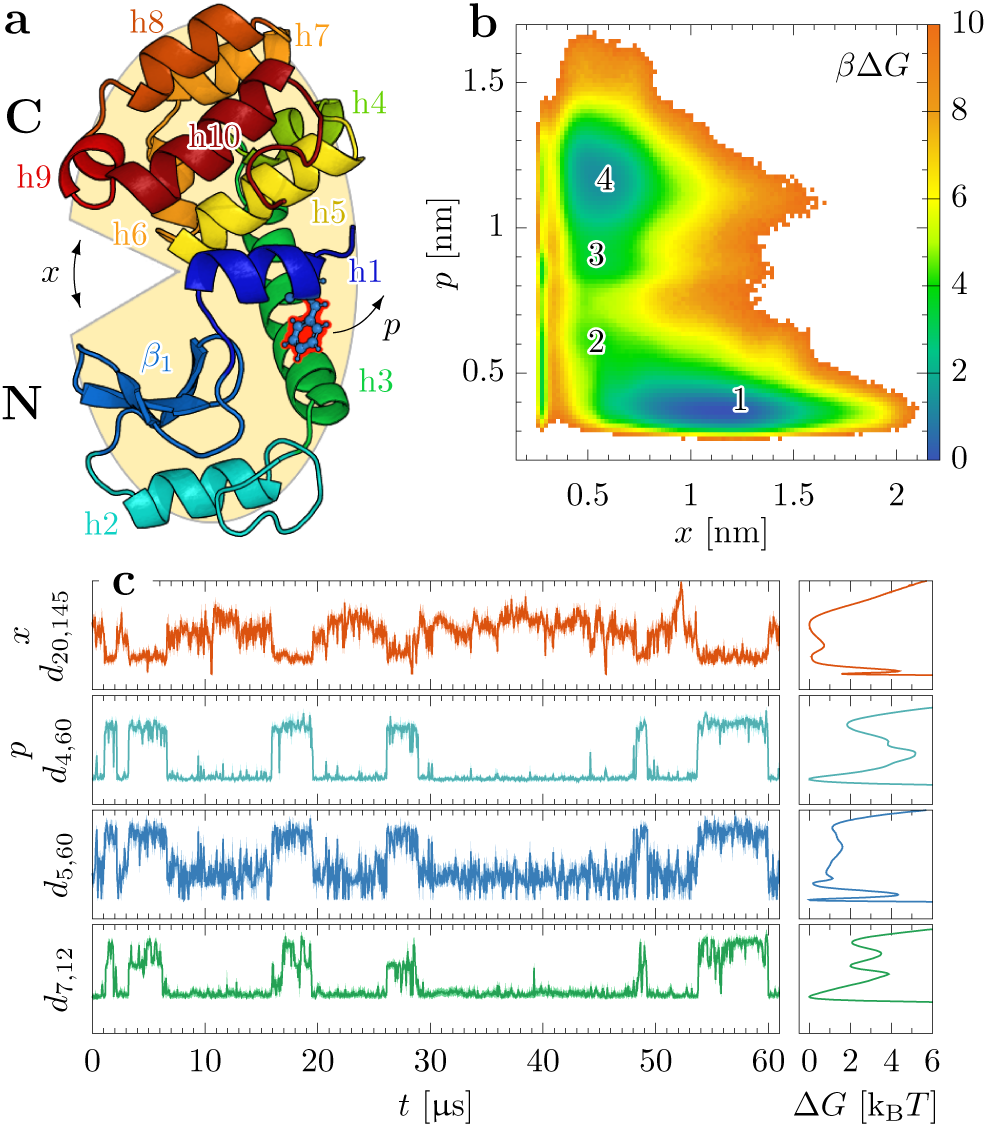
(a) Structure of the N- and C-domain of T4L, indicating the opening coordinate *x* of the “mouth” region and the locking coordinate *p* that describes the motion of key residue Phe4 (red) in the “hinge” region. (b) Two-dimensional free energy landscape Δ*G*(*x, p*) (in units of *k*_B_*T*), indicating four conformational states. (c) MD time traces of various distance coordinates describing the open-closed transition and their corresponding free energy profiles.

Combining a correlation analysis of contact features^32^ with a careful inspection of individual transitions, we show that the open-closed transition of T4L amounts to a cooperative process, where a large number of inter-atomic distances from the hinge to the mouth region act in a simultaneous manner. In particular, we find that the hydrophobic core of the hinge region constitutes a flexible transmission network, which represents the molecular machinery at the heart of the open-closed transition.

## RESULTS AND DISCUSSION

### Two-dimensional model: the locking mechanism

The open-closed motion of T4L is usually described by the mouth opening width *x* (Fig. 1a), which can be represented, e.g., by the distance *d*_20,145_ between residues 20 and 145 of the N- and C-domain, respectively. Employing the MD trajectory by Ernst et al.,^29^ Fig. 1c reveals that the open-closed transitions of *x*(*t*) occur on a microsecond timescale, while the transition path time lasts only nanoseconds. From the data we estimate equilibrium populations of 70 % (open) and 30 % (closed) and mean waiting times of *τ*_o*→*c_ *∼* 4 µs and *τ*_c*→*o_ ∼ 2 µs, which is in excellent agreement with recent experimental results.^27^ The corresponding free energy profile Δ*G*(*x*), however, exhibits a rather low energy barrier (Δ*G ∼* 1*k*_B_*T*). ≠ Adopting the standard expression for the transition rate, 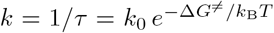, it is obvious that the rate is almost solely caused by the prefactor *k*_0_.^33^ This indicates that the opening width *x* does not reveal the underlying mechanism of the process and is therefore not a suitable one-dimensional reaction coordinate.

Indeed, when we employed targeted MD simulations to enforce the open-close motion by pulling along *x*, the two-state behavior of T4L could not be recovered.^29^ Rather, the transition was found to be triggered by a locking mechanism, by which the side chain of Phe4 changes from a solvent-exposed to a hydrophobically-buried state. Describing the motion of Phe4 via the hydrophobic contact distance between Phe4 and Lys60, *d*_4,60_ ≡*p*, this coordinate discriminates between locked (*p* 0.7, hydrophobically buried) and free (*p* 0.7, solvent exposed) states.

Figures 1b,c show the free energy landscape Δ*G*(*x, p*) and the time traces *x*(*t*), *p*(*t*) of the resulting two-dimensional model of T4L. The landscape exhibits four metastable conformational states labeled by numbers **1** - **4**, including the prominent open locked state **1** and closed free state **4**, as well as two sparsely populated intermediate closed states **2** and **3**.

The locking mechanism is found to stabilize the open and closed states and can therefore be considered as the origin of the bistable behavior of T4L. Demonstrating the coupling of opening coordinate *x* and locking coordinate *p*, the free energy landscape Δ*G*(*x, p*) also provides first mechanistic insights of the conformational transition.^29^ In particular Δ*G*(*x, p*) indicates that most transitions proceed along the route **1***→***2***→***3***→***4** and back along **4**→ **3** →**2** →**1**. On the other hand, the two-dimensional model of T4L apparently still does not include some important aspects of the dynamics, because the corresponding highest free energy barrier is still relatively small (∼ 5 *k*_B_*T*). As we show in this work, the missing link is the long-range propagation of conformational change, which couples the (un)locking of Phe4 in the hinge region to the opening/closing of the mouth region.

### Correlation analysis: cooperative dynamics

To describe the mechanistic coupling between the mouth and hinge regions in T4L, we want to identify internal coordinates that change significantly during the open-closed transition. Following Ernst et al.,^29^ we consider interresidue contact distances and side-chain *χ*_1_ dihedral angles, as they report on near-order interactions, from which long-distance changes result as a consequence. Assuming that a contact is formed if the distance *d*_*ij*_ between the closest non-hydrogen atoms of residues *i* and *j* is shorter than 4.5 °A,^34^ we identified 844 interresidue contacts that are formed at least 1 % of the simulation time. Moreover, we focused on the first side-chain dihedral angle of each residue, resulting in 131 coordinates *χ*_*n*_ (Ala, Gly and Pro do not exhibit or change this angle). To study which of these coordinates {*x*_*α*_} = {*d*_*ij*_, *χ*_*n*_ }are interrelated to each other, we calculated the linear correlation matrix

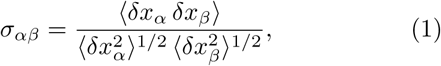

where *δx*_*α*_ = *x*_*α*_ − ⟨*x*_*α*_ ⟩and ⟨… ⟩ denotes the time average over the MD data. Aiming to discriminate collective motions underlying functional dynamics from uncorrelated motions, we wish to rearrange this matrix in an approximately block-diagonal form. Employing the Python package MoSAIC,^32^ this is achieved via a community detection technique called Leiden clustering.^35^

The resulting block-diagonalized correlation matrix in Fig. 2a shows a main cluster (i.e., the first block) that contains 85 strongly correlated coordinates, which account for the open-closed transition (see below). Moreover, we obtain six smaller clusters describing local motions that are not related to the open-closed motion (Fig. S1), and the remaining 530 clusters that exhibit only minor correlation and are therefore considered as noise (see Table S1 for a list of all clusters and coordinates). The latter finding is associated with the fact that in T4L intraprotein contacts are quite stable and only fluctuate around their mean distance, while contacts on the protein surface frequently form and break and hence fluctuate randomly. That is, of the 976 considered internal coordinates only 85 (82 contact distances and 3 dihedral angles) are in fact associated with the open-closed transition of T4L.

**FIG. 2.**
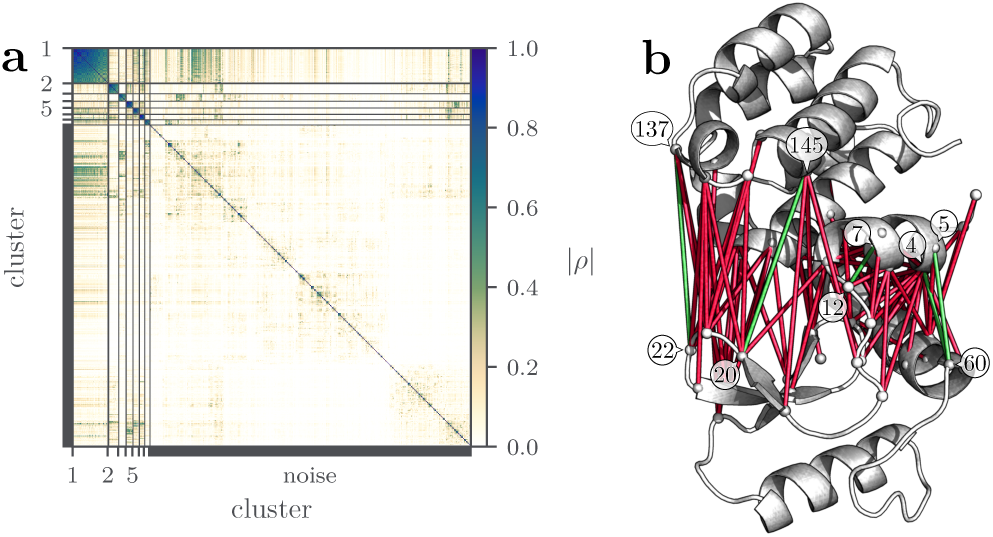
(a) Block-diagonal form of the correlation matrix built from 844 contact distances and 131 side-chain dihedral angles of T4L. The first block (cluster 1) contains 85 highly correlated coordinates that account for the open-closed transition. (b) Illustration of the 82 relevant contact distances in red, as well as in green the important distances *d*_4,60_, *d*_5,60_, *d*_22,137_, *d*_20,145_ describing the opening of the mouth, and *d*_7,12_ accounting for the (de)stabilization of helix 1.

Displaying these 82 highly correlated distances in the molecular structure of T4L, Fig. 2b shows that the coordinates span the space connecting the mouth and hinge regions. They include 36 mouth-opening coordinates and 15 coordinates involving Phe4, while the remaining coordinates are mostly located in the hinge region. As demonstrated by the time evolution of some important examples (Fig. 1c), these coordinates follow the microsecond open-closed transitions of T4L. Hence, the above correlation analysis indicates that the open-closed transition is mediated via the 82 highly correlated contact distances that connect the hinge and mouth regions of T4L.

While the overall time evolutions of the coordinates in Fig. 1c are clearly correlated, the various contact distances are seen to differ in their fluctuations in the open and closed state. E.g., opening coordinate *x* fluctuates considerably in the open state and only little in the closed state, while it is the other way round for the locking coordinate *p*. This is because amino acid side-chains that are in contact generally fluctuate significantly less than in the case of a broken contact. Although the fast fluctuations of different contacts are not necessarily correlated, the open-close transition may only take place if the fluctuations of the various contacts by chance cause the simultaneous switching of all relevant interactions. Hence we propose that a successful transition necessitates a concerted or cooperative action of all involved coordinates of the intramolecular contact network of T4L.

### Single event analysis: essential coordinates

To facilitate a concise discussion of the structural changes associated with the open-closed transition, we need to further reduce the number of coordinates. In a first step, we disregard all contacts that do not change during the open-closed transition, but only fluctuate in either the contact-formed or contact-broken state. This leaves us with 18 hydrophobic contacts, 5 hydrogen bonds, and 4 salt bridges (Table S2). While hydrophobic contact changes typically involve complex rearrangements of various atoms that are difficult to grasp (see below), changes of hydrogen bonds and salt bridges can often be directly linked to overall structural changes and are therefore considered first.

Most prominently, we identify three salt bridges that change during the open-closed transition. That is, in the hinge region the salt bridge between Glu5 and Lys60 opens, while in the mouth region the salt bridges Asp20-Arg145 and Glu22-Arg137 close. The first salt bridge described by *d*_5,60_ complements the previously defined locking coordinate *p* ≡*d*_4,60_, while the second salt bridge corresponds to the above discussed opening coordinate *x* ≡*d*_20,145_. Ultimately, the third salt bridge is associated with the contact distance *d*_22,137_ describing the opening and closing of the tip of the mouth, see Fig. 2b. Apart from these fairly obvious interactions, we find one more salt bridge (between Arg8 and Glu64) and several hydrogen bonds that are located in the mouth region (described by *d*_21,105_ and *d*_22,141_) and in the hinge region (*d*_2,64_, *d*_7,11_, and *d*_7,12_).

As a simple means to study the mutual relation of these coordinates, we considered their time evolution during individual open ↔ closed transitions of the trajectory (Fig. S2). Interestingly, we find that the salt bridge *d*_8,64_ and the hydrogen bond *d*_2,64_ compete with each other, facilitating a seesaw-like motion of helix 1 with respect to helix 3 (Fig. S3). Moreover, we note that hydrogen bond distances *d*_21,105_ and *d*_22,141_ show quite similar behavior as the main salt bridges and are therefore redundant. On the other hand, distance *d*_7,12_ (or alternatively *d*_7,11_) is found to exhibit diverse behavior, as it accounts for the (de)stabilization of helix 1 (see below). From the three side-chain dihedral angles of cluster 1 (*χ*_4_, *χ*_70_, and *χ*_104_), *χ*_4_ is most important, as it is found to directly account for the re-orientation of Phe4. While *χ*_104_ exhibits a similar behavior over time as *χ*_4_, it shows clearly more fluctuations whose structural implications are resolved below. *χ*_70_ shows hardly additional information and is therefore neglected. Put together, this leaves us with coordinates *d*_5,60_, *d*_4,60_, *χ*_4_ and *d*_7,11_ reporting on the (un-)locking of Phe4 and the subsequent rearrangement of helix 1, as well as *d*_20,145_ and *d*_22,137_ describing the opening and closing of the mouth. Illustrating the mechanism of the transition, these coordinates can be considered as essential internal coordinates.^30^

To illustrate to what extent these coordinates from the hinge to the mouth region indeed move simultaneously, we zoom into the time evolution of the system during a representative open →closed transition. Assigning the start of the transition to time *t* = 0, Fig. 3a reveals that distance *d*_5,60_ changes first, before right afterwards coordinates *d*_4,60_, *d*_22,137_ and *d*_20,145_ change virtually simultaneously. A few nanosecond later *χ*_4_ responds, followed by *d*_7,12_ about 10 ns later. Of particular interest is the simultaneous response of distances *d*_4,60_ (hinge) and *d*_22,137_ (mouth), which are far apart (about 2.5 nm) from each other (see Fig. 2b). This synchronism indeed seems to indicate a direct mechanical coupling of the two protein sections.

**FIG. 3.**
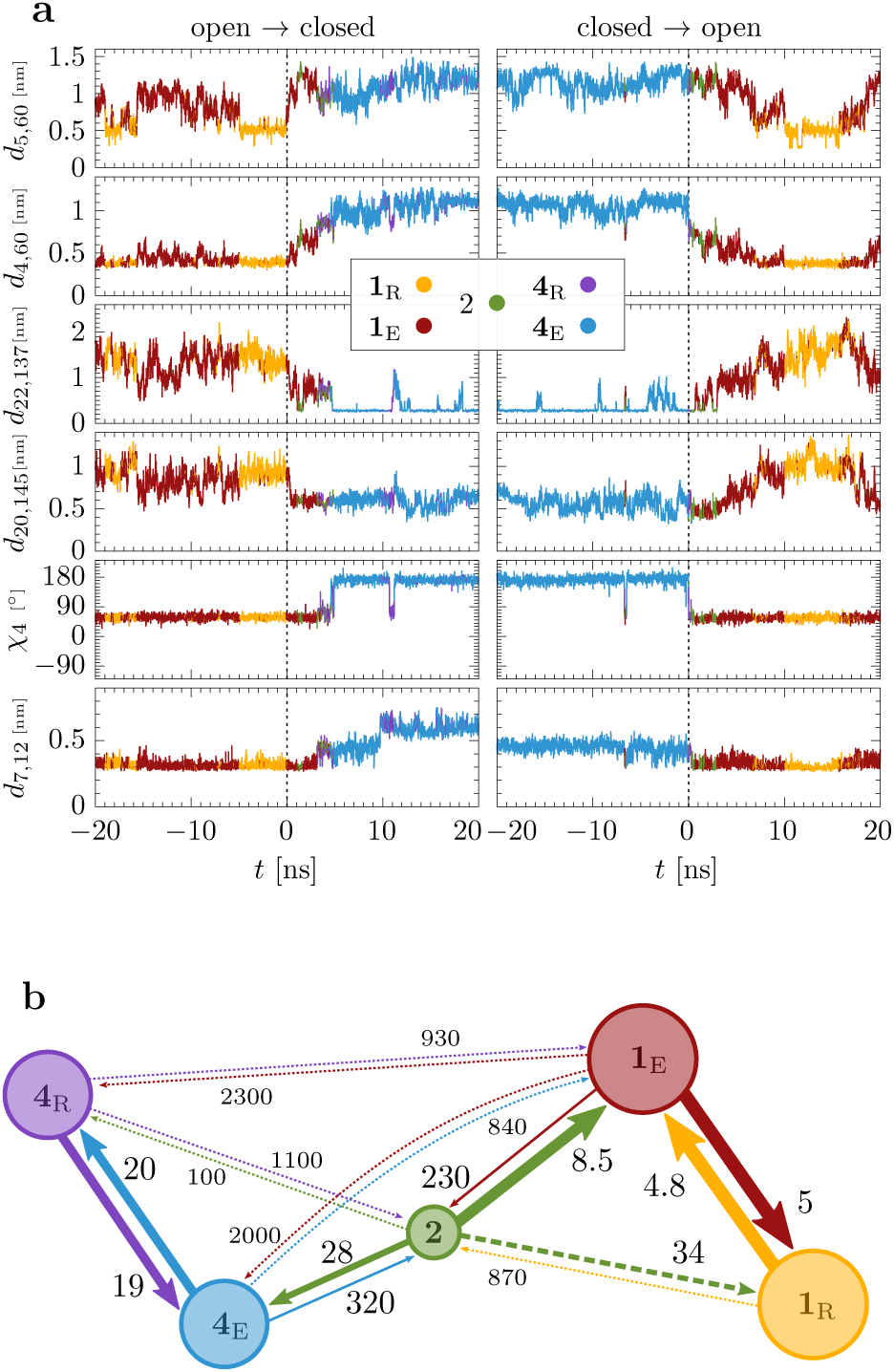
(a) Time evolution of some key coordinates of T4L during a representative open*→*closed (left, occurring at 3.2 µs in Fig. 1) and closed*→*open (right, at 16.6 µs) transition, respectively. The color encodes the instantaneous conformational state of the system revealed by density-based clustering. (b) Scheme of the resulting Markov state model indicating the main transition times (in units of ns).

### Structural characterization: transmission network

To explain the above findings, Fig. 4a depicts the overall conformational rearrangement of T4L during the open-closed transition. While the N- and C-terminal protein domains perform a mirrored rotational motion, helix 1 executes a piston-like move into the opposite direction. This structural coupling between N- and C-terminal domains as well as with helix 1 is facilitated by a network of persistent salt bridges indicated in Figs. 4b and S3. That is, helix 1 is connected to the C-terminal domain via several salt bridges between the two negatively charged residues Asp10 and Glu11 in helix 1 and the two positively charged residues Arg145 and Arg148 in helix 10 of the C-terminal domain. Along helix 3, we find a salt bridge between Asp72 and Arg76. Furthermore, helix 3 is connected to the N-terminal domain via Arg52-Glu62, and to the C-terminal domain via Arg80-Glu108. All these electrostatic connections serve as structural tethers that provide a balance between stability and flexibility, which we consider as a prerequisite for a bistable system. In Fig. 4b, panels (1) to (5) comprise the individual local sites mediating the conformational rearrangement from the protein open to the closed state. Moreover, the analysis suggests the intramolecular coordinates shown in Fig. 3a, which are indicative of the local structural changes. The structural evolution is illustrated in real time in the Supplementary Movie S1.

**FIG. 4.**
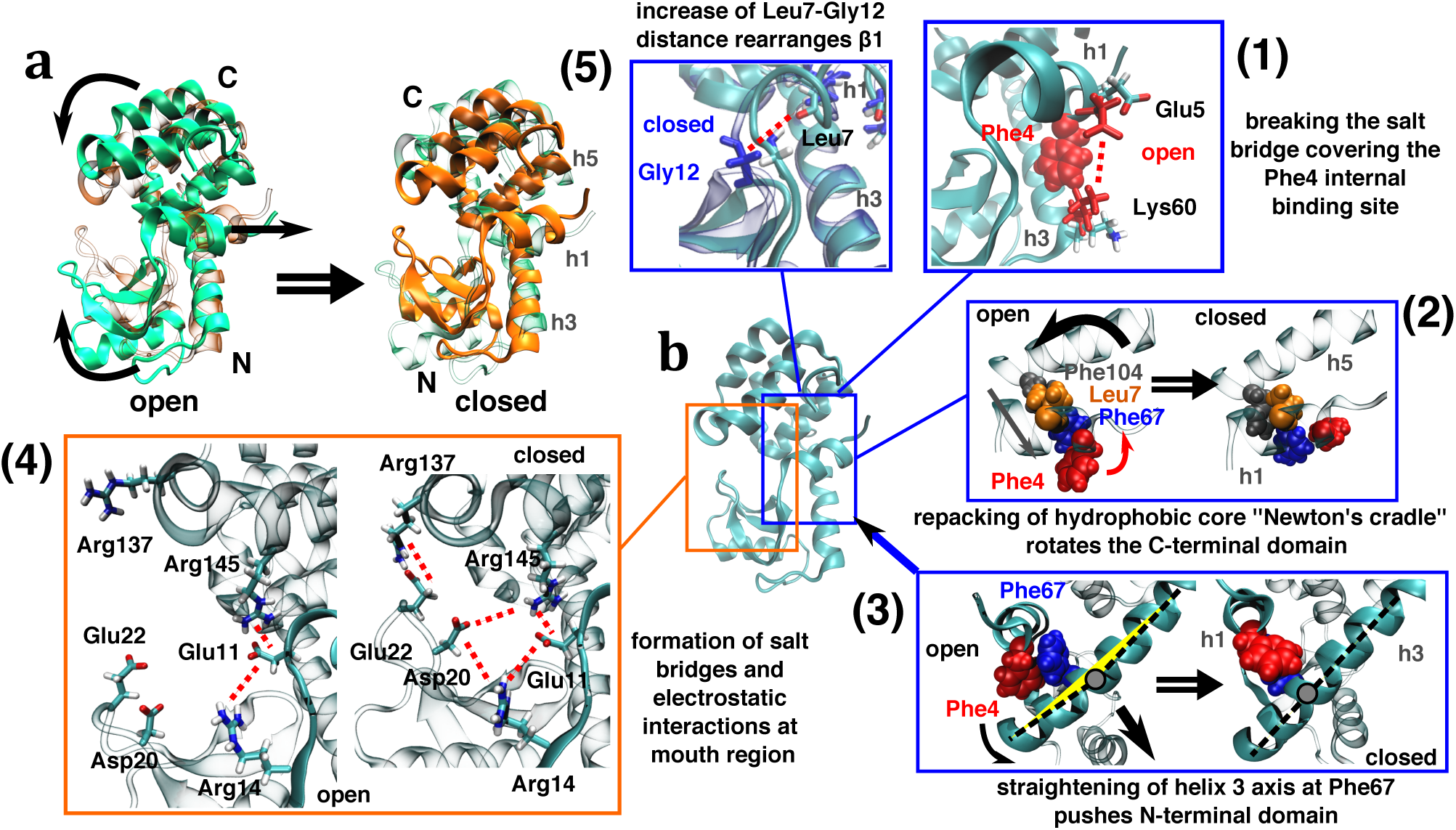
Structural details of the open-closed mechanism in T4L, with black arrows indicating the direction of backbone motions. (a) illustrates the overall picture, (b) accounts for the individual steps of the conformational rearrangement, see the text.

1. To allow the exit of Phe4 from its hydrophobic pocket, first a salt bridge between Glu5 and Lys60 needs to open. This motion is naturally described by the contact distance *d*_5,60_. The subsequent outward move of Phe4 can be monitored by the locking coordinate *d*_4,60_.
2. The outward motion of Phe4 is transferred via Phe67, Leu7 to Phe104 and its adjacent helix 5. That is, Phe4, Phe67, Leu7 and Phe104 form a hydrophobic core that causes a rolling motion of helix 5 over helix 1 similar to two cogwheels in contact. As helix 5 represents an integral part of the C-terminal domain, this motion of helix 5 rotates the C-terminal domain as shown in Fig. 4a. At the end of the rotation, Phe104 on the opposite end of helix 1 reacts via a change of its first side-chain dihedral angle *χ*_104_.
3. At the same time, the conformational change of the Phe67 side chain is transferred farther to the backbone of helix 3, which causes a straightening of the helix. This straightening acts on the N-terminal domain via the backbone and side chain packings, causing the forward movement of the N-terminal domain. In combination, the above described motions of the C- and the N-terminal domains lead to the closing of the mouth of T4L.
4. The closed conformation is stabilized via a salt bridge at the tip of the mouth between Glu22 and Arg137, which can be followed via distance *d*_22,137_, and an electrostatic interaction complex involving Glu11, Arg14, Asp20 and Arg145 that can be monitored by *d*_20,145_ in the middle of the mouth. Both salt bridge centers impose structural near-order on the mouth region. Since these coordinates respond simultaneously with the locking coordinate *d*_4,60_ (which indicates that the Phe4 side chain passes over the side chain of Phe67), the complex conformational rearrangements described in steps (2) to (4) occur in a concerted manner. Subsequently, the Phe4 side chain rotates into a new conformation and becomes completely exposed to the solvent. This can be monitored by the side-chain dihedral angle *χ*_4_.
5. Finally, a destabilization of the last winding of helix 1 leads to a rupture of a backbone hydrogen bond between Leu7 and Gly12, resulting in an increase in the distance *d*_7,12_. As a consequence, the motion of Gly12 causes a change in the position of the *β*_1_ strand such that Arg14 is rotated away from the electrostatic complex, thus weakening its coupling with Glu11 and Asp20.

To go back to the open state, the above steps simply occur in reverse order. Here the crucial issue is the locking of Phe4, i.e., its move from the solvent-exposed to a hydrophobically-buried state, which is indicated by coordinates *d*_7,12_, *χ*_4_ and *p*.

In summary, we have shown that the hydrophobic core (Phe4, Phe67, Leu7 and Phe104) mediating a cogwheel-like motion of helices 1 and 5 and the straightening of helix 3 constitutes a flexible transmission network, which represents the molecular machinery underlying the open-closed transition. While common motions of Phe4, Phe67 and Phe104 seemed plausible from the comparison of various wild-type and mutant crystal structures of T4L,^24,25^ here we show that these residues together with Leu7 in fact perform a concerted motion. Since helices 3 and 5 are rigidly connected to the N- and C-terminal domains, respectively, the motion of the transmission network in the hinge region is directly and instantaneously coupled to the mouth region of T4L. This long-range or allosteric coupling is the origin of the concerted action of the 85 distances connecting the mouth and the hinge region of T4L (Fig. 2b).

### State model description: heterogeneity of pathways

The finding of several steps in the cooperative mechanism raises the question to what extent these steps are associated with well-defined intermediate states of the process. To this end, we employed the six coordinates (*x, p, d*_5,60_, *d*_22,137_, *χ*_4_, *d*_7,12_) discussed in Fig. 3a and performed robust density-based clustering^36^ in this space (see Material and methods for details). In line with the nomenclature of the previous four-state model (Fig. 1c), the resulting five conformational states are referred to as states **1**_R_ (with a population of 33.5 %), **1**_E_ (35.5 %), **2** (1.3 %), **4**_E_ (15.1 %) and **4**_R_ (14.5 %), see Fig. 3b. Compared to the previous model that relied on coordinates (*x, p*) only, notably the two main states **1** (open) and **4** (closed) each split up into two similarly populated substates, indexed by “R” (for relaxed) and “E” (for excited). In both cases, the sub-state E represents the “pole position” to start the open*↔*closed transition, while the sub-state R accounts for the fully relaxed equilibrium state. Displaying the probability distributions of all six coordinates in each state, Fig. S4 reveals that the splitting of state **1** is caused by coordinate *d*_5,60_, while the splitting of state **4** is caused by coordinate *d*_7,12_. Although the intermediate state **2** is weakly populated and short-lived as in the previous model, it turns out to be mechanistically relevant (see below). In contrast, the weakly populated previous intermediate state **3** is now a part of state **4**_E_. The spatial extension and the connectivity of the five states are well characterized by the two-dimensional free energy surfaces along the above defined six coordinates shown in Fig. S5.

To relate these conformational states to steps (1) - in Fig. 4b, we consider the resulting state trajectory during the representative open → closed transition shown in Fig. 3a. As suggested above, we find that transition **1**_R_ → **1**_E_ accounts for the initial unlocking of Phe4 (Fig. 4b1). The subsequent main cooperative process (Figs. 4b2 - 4), on the other hand, is associated with two transitions, **1**_E_ → **2** and **2** → **4**_E_. State **2** represents the first stabilized closed structure, that is, it succeeded to stabilize one of the frequent attempts to close the mouth along opening coordinate *x*. Moreover, we find that helix 3 is already straightened in this state, reflecting step (3). Only by reaching state **4**_E_, though, the cogwheel-like motion of the hydrophobic core and helices 3 and 5 (steps (2) and (3)) can be completed, with Phe4 moving into the solvent. Finally, transition **4**_E_ → **4**_R_ describes the stabilization of the closed state (step (5)), which typically takes longer than the nanosecond timescale shown in Fig. 3a.

To illustrate the heterogeneity of the open↔closed transitions and discuss the possible pathways of the reaction, Fig. S2 shows additional transition events apart from the one shown in Fig. 3a. Moreover, we calculated the resulting transition matrix *T* (*τ*_lag_) containing the probabilities *T*_*ij*_ that the system jumps from state *i* to *j* within lag time *τ*_lag_. To distinguish completed transitions from transient fluctuations, dynamical coring^37^ was applied which requires that the trajectory spends a minimum time *τ*_cor_ in the new state for the transition to be counted (Fig. S6). Using *τ*_cor_ = *τ*_lag_ = 0.7 ns, the resulting transition times *t*_*ij*_ = *τ*_lag_*/T*_*ij*_ of the state model are shown in Fig. 3b. We note that the state doublets (**1**_R_, **1**_E_) and (**4**_R_, **4**_E_) interconvert frequently with 5 and 20 ns transition time, respectively. Apart from the main pathway **1**_R_→ **1**_E_ → **2** → **4**_E_ → **4**_R_ discussed above, we find various shortcuts of the reaction that, e.g., omit state **1**_E_ (by going directly from **1**_R_ to **2**) or omit state **2** (by going directly from **1**_E_ to **4**_E_), or omit state **4**_E_ (by going directly from **2** to **4**_R_).

When we attempt to construct a Markov state model from the transition matrix, we find that its predictions of the state-to-state waiting times are only in qualitative agreement with the corresponding MD results, i.e., they deviate from MD results by a factor of ∼ 2 (Fig. S6). This reflects the fact that during a successful cooperative open-close reaction the various state-to-state transitions occur almost simultaneously (Fig. 3a), i.e., they are not independent from each other.^38^ This is opposed to the timescale separation assumed by a Markov state model, where the lag time (representing the randomization time of the fast intrastate dynamics) should be significantly shorter than the transition times of the model.

## CONCLUSIONS

To investigate the microscopic mechanism underlying the open-closed transition of T4L, we have combined a correlation analysis of contact features (Fig. 2) with a study of their time evolution (Fig. 3) and a careful visualization of the conformational changes (Fig. 4 and Supplementary Movie S1). In particular, we have identified a set of essential internal coordinates (Fig. 3) that monitor various aspects of the transition. The allosteric mechanism in T4L is found to consist of three ingredients.

i. To start the open-closed transition, a salt bridge between Glu5 and Lys60 needs to open, in order to allow the exit of Phe4 from its hydrophobic pocket (Fig. 4b1). The finalization of the overall transition requires a rearrangement of salt bridges in the mouth region (Fig. 4b4) and a hydrogen bond of helix 1 (Fig. 4b5), which both stabilize the closed state. In the case of the converse reaction, first the latter three interactions are reversed, while the final formation of the Glu5-Lys60 salt bridge stabilizes the closed state. That is, both open and closed states are stabilized by specific interactions, which initially need to be overcome to get the system in a pole position for the desired transition.
ii. Following this initiation, the hydrophobic core (Phe4, Phe67, Leu7 and Phe104) of the hinge region mediates a cogwheel-like motion of helices 1 and 5 (Fig. 4b2) as well as the straightening of helix 3 (Fig. 4b3). In this way, the hydrophobic core constitute a flexible transmission network, which represents the molecular machine driving the open-closed transition.
iii. Since helices 3 and 5 are tightly connected to the N- and C-terminal domain, respectively, the motion of the transmission network in the hinge region is directly and instantaneously coupled to the mouth region of T4L. Hence the small local changes of the hydrophobic core are passed on via rigid contacts and the protein backbone to distant sites.

We anticipate that these three components of the process are ubiquitous for allosteric transitions of bistable proteins. For example, a repacking of a hydrophobic core as well as switching of polar interaction partners was reported for the nitrogen regulatory protein C.^17^ Moreover, the above described lever-arm effect is reminiscent of structural changes in G protein-coupled receptors, where small changes in the ligand binding site are structurally amplified to large-scale protein surface changes at the G protein binding site.^39,40^ A similar effect was also reported for the microbial rhodopsin bacteriorhodopsin, where the retinal cofactor pushing against Trp182 causes a large-scale outward motion of helix F,^41^ and for the connection between ATP hydrolysis and protein conformational changes in heat shock protein 90.^20^ The interplay between rigid secondary elements (such as *α*-helices and *β*-sheets) and flexible protein sections (such as loops or linkers) to mediate structural rearrangements was also discussed by Nussinov and Thirumalai.^9,42^

The simultaneous involvement of residues of the N- and C-terminal domains as well as of helix 1 demonstrates a high cooperativity of the open-closed transition. This finding is in line with recent unfolding experiments on T4L,^26^ which revealed that helix 1 is responsible for the high degree of cooperativity within the protein. By coupling the two domains, helix 1 gives rise to an all-or-none, two-state unfolding behavior. It is interesting to note that the cooperativity of the (un)folding of T4L seems to transfer to the cooperativity of its functional dynamics. This insight could allow us to employ –at least to some extent– the well-established theories and concepts of protein folding to the less understood theoretical description of allosteric transitions. In this context we note that recent time-resolved experimental and computational studies of a photoswitchable PDZ2 domain indicated that allosteric communication shares some properties with multistate downhill folding.^8,19^ Similar to the various degrees of cooperativity in protein folding,^43^ allosteric transitions may as well show a variety of mechanisms.

Using the identified essential internal coordinates, density-based clustering revealed five conformational states (Fig. 3b), which showed a splitting of the two main open and closed protein states reflecting their stabilization described in (i). Apart from this and a weakly populated optional intermediate state, however, the resulting Markov state model provided only little insight in the cooperative open-close transition. This is because in a cooperative process the various motions occur almost simultaneously (Fig. 3a) and are therefore not independent from each other, which is opposed to the timescale separation assumed by a Markov model.

Cooperative processes such as two-state protein folding, on the other hand, have been successfully described by Ising-type models.^44,45^ Here we consider an Ising model below the critical temperature, where local couplings between two-state entities lead to the emergence of well-defined global orientations of the system. Applied to the functional dynamics of a bistable protein, we associate the local two-state systems with the side-chain contacts, their next-neighbor couplings with interactions with residues in side-chain or backbone proximity, and the resulting global orientations with the open and closed protein conformations. In direct analogy to the system’s stabilization in a single global orientation by an external field, in allostery a protein state can be stabilized via the (un)binding of a ligand.

The Ising model approach explains the cooperativity of the global open-closed transition via the interaction of local fluctuations.^46^ While all local contacts fluctuate on their own, a successful conformational transition requires the simultaneous flipping of all contacts at the same time. In other words, the system needs to pass through a narrow region in phase space, which constitutes a rare event.^47^ In particular, the need of a simultaneous change of numerous contacts explains the long waiting times despite small barriers along individual coordinates (Fig. 1). A similar combination of small barriers and transitions requiring maximal cooperativity between amino acids was found for the folding of small proteins.^48^

In conclusion, we have shown for a case study of T4L that the combination of extensive MD simulations, correlation-based dimensionality reduction, detailed inspection of the structural changes, and models from protein folding theory may provide a microscopic understanding of the allosteric mechanism in unprecedented detail. As in an Ising model, global conformational rearrangements and allosteric communication over long distances are found to emerge from local fluctuations and short-distance couplings.

## MATERIAL AND METHODS

### MD simulations

This work is based on the 61 µs- long all-atom MD trajectory of Ernst et al.,^29^ which used the GROMACS v2016 software package^49^ and the Amber99*ILDN force field.^50–52^

### Definition of coordinates

The Leiden algorithm^35^ is a community detection algorithm which identifies subparts of strongly interacting nodes in graphs. For the identification of collective motion in T4L we constructed a similarity-graph in which the nodes represent the coordinates and the edges indicate how correlated they are. For this graph, the Leiden algorithm maximizes an objective function, indicating to what extent the graph exhibits a clusters-like topology. Here we employed the constant Potts model as the objective function

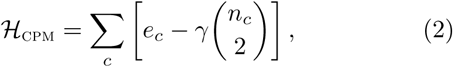

where *e*_*c*_ denotes the sum of edge weights and *n*_*c*_ the number of nodes in cluster *c. γ* is a resolution parameter that governs the degree to which coordinates within a cluster must be correlated. Setting *γ* = 0.5, seven larger clusters were found, while the majority of the coordinates are mostly uncorrelated to the rest of the system. Cluster 1 corresponds to 85 coordinates, listed in Table S1, that correlate highly with the open-closed transition.

### Characterization of metastable states

Robust density-based clustering^36^ identifies states based on a local free energy estimate which is calculated for each trajectory point by counting the number of neighbors found inside a *d*-dimensional hypersphere of fixed radius *R* centered at the point. We applied this method to the coordinates *x* = *d*_20,145_, *p* = *d*_4,60_, *d*_5,60_, *d*_22,137_, *χ*_4_ and *d*_7,12_. To ensure that all six coordinates span similar value ranges the angle *χ*_4_ was divided by 180. Using a clustering radius of *R* = 0.034 and a minimal population *P*_min_ = 100 resulted in 24 state clusters. Of those 24 states, only the first 9 states have populations *>* 1 %. By inspecting the location of these states in the six-dimensional space, we were able to associate them with one of the 5 coarse-grained states **1**_R_ (microstates 2 and 5), **1**_E_ (microstate 1), **2** (microstate 9), **4**_R_ (microstates 4 and 8) and **4**_E_ (microstates 3, 6 and 7). The remaining 15 microstates were assigned to the closest coarse-grained state.

### Markov state model

We first applied dynamical coring to improve the state definition.^37^ Here, short-living fluctuation are removed from the dynamics by requiring for valid transitions that the trajectory spends at least a minimum time *τ*_cor_ in the new state. To determine *τ*_cor_, we considered the probability *P*_stay,*i*_(*t*) to stay in state *i* which should decay like a single exponential function. Short-lived fluctuation induce a fast initial decay, which can be removed by using *τ*_cor_ = 700 ps. To test the reliability of the model, we generated a Markov chain Monte Carlo trajectory and compared the average waiting times of MD and Markov model, i.e., the average times *t*_wait,i*→*j_ needed to reach state *j* after entering state *i* (Fig. S6).

## Supporting information

Supplementary Informations

## Acknowledgments

The authors thank Peter Hamm, Thorsten Hugel, Peter Bolhuis, Daniel Nagel, and Simon Stitzinger for fruitful discussions. This work has been supported by the Deutsche Forschungsgemeinschaft (DFG) via the Research Unit FOR 5099 “Reducing complexity of nonequilibrium” (project No. 431945604). The authors acknowledge support by the bwUniCluster computing initiative, the High Performance and Cloud Computing Group at the Zentrum für Datenverarbeitung of the University of Tübingen, the state of Baden-Württemberg through bwHPC and the DFG through grant No. INST 37/935-1 FUGG.

