## Supplementary Informations for "Cooperative protein allosteric transition mediated by a fluctuating transmission network"

Matthias Post, Benjamin Lickert, Georg Diez, Steffen Wolf,* and Gerhard Stock†
*Biomolecular Dynamics, Institute of Physics, Albert Ludwigs University, 79104 Freiburg, Germany.*
(Dated: April 28, 2022)

### I. SUPPLEMENTARY TABLES

TABLE S1: Inter-residue distances and first side-chain dihedral angles within cluster 1 found by the Leiden algorithm [1], sorted by their average correlation within the cluster. Comparing various correlation measures and several clustering algorithms, a recent study [2] has shown that this result is quite robust and depends only little on details of the clustering.

| Cluster | Coordinates |
|---|---|
| 1 | $d_{4,15}$, $d_{4,28}$, $d_{4,60}$, $d_{4,63}$, $d_{23,141}$, $d_{4,13}$, $d_{4,61}$, $d_{22,137}$, $d_{20,142}$, $d_{21,141}$, $d_{20,141}$, $d_{22,141}$, $d_{21,137}$, $d_{24,137}$, $d_{21,143}$, $d_{4,29}$, $d_{21,138}$, $d_{21,142}$, $d_{32,106}$, $d_{24,141}$, $d_{8,67}$, $d_{4,72}$, $d_{2,64}$, $d_{30,145}$, $d_{32,105}$, $d_{4,64}$, $d_{8,68}$, $d_{67,100}$, $d_{7,71}$, $d_{32,107}$, $d_{1,64}$, $d_{35,106}$, $d_{7,13}$, $d_{22,142}$, $d_{20,145}$, $d_{8,104}$, $d_{4,75}$, $d_{20,105}$, $d_{35,109}$, $d_{7,12}$, $d_{24,106}$, $d_{5,60}$, $d_{7,75}$, $d_{4,71}$, $d_{4,69}$, $d_{35,107}$, $d_{24,105}$, $d_{9,13}$, $d_{8,64}$, $d_{18,145}$, $d_{8,13}$, $d_{7,74}$, $d_{8,12}$, $d_{24,107}$, $d_{4,68}$, $d_{32,104}$, $\chi_4$, $d_{5,64}$, $d_{3,67}$, $d_{24,145}$, $\chi_{101}$, $d_{11,20}$, $d_{11,26}$, $d_{10,29}$, $d_{22,106}$, $d_{11,28}$, $d_{7,11}$, $d_{14,145}$, $d_{67,101}$, $d_{29,145}$, $d_{8,15}$, $d_{2,67}$, $d_{4,9}$, $d_{65,70}$, $d_{22,105}$, $d_{21,105}$, $d_{11,30}$, $d_{66,71}$, $d_{31,104}$, $d_{66,70}$, $d_{30,104}$, $d_{21,106}$, $d_{5,67}$, $\chi_{70}$, $d_{75,88}$ |
| 2 | $d_{33,46}$, $d_{17,33}$, $d_{33,45}$, $d_{33,43}$, $d_{33,42}$, $d_{27,31}$, $d_{33,41}$, $d_{27,33}$, $d_{25,33}$, $d_{18,33}$, $d_{27,32}$, $d_{33,49}$, $d_{32,45}$, $d_{31,66}$, $d_{32,49}$, $d_{33,50}$, $d_{31,50}$, $d_{19,24}$, $d_{25,31}$, $d_{31,46}$, $d_{31,58}$, $d_{25,32}$, $d_{18,26}$ |
| 3 | $d_{6,98}$, $d_{10,101}$, $d_{6,100}$, $d_{6,97}$, $d_{9,161}$, $d_{6,94}$, $d_{9,160}$, $d_{3,100}$, $d_{9,158}$, $d_{10,105}$, $d_{6,101}$, $d_{10,149}$, $d_{10,145}$, $d_{9,148}$, $d_{3,96}$, $d_{6,152}$ |
| 4 | $d_{34,39}$, $d_{34,42}$, $d_{34,38}$, $d_{25,34}$, $d_{36,42}$, $d_{19,34}$, $d_{36,45}$, $d_{26,34}$, $d_{24,34}$, $d_{34,41}$, $d_{37,41}$, $d_{23,34}$, $d_{17,34}$, $d_{35,45}$ |
| 5 | $d_{18,22}$, $d_{20,24}$, $d_{14,22}$, $d_{14,21}$, $d_{20,25}$, $d_{22,26}$, $d_{14,20}$, $d_{20,32}$, $d_{22,30}$, $d_{20,26}$, $\chi_{20}$, $d_{11,22}$, $d_{20,30}$ |
| 6 | $d_{18,30}$, $d_{18,29}$, $d_{18,28}$, $d_{18,27}$, $\chi_{18}$, $d_{18,31}$, $d_{12,18}$, $d_{13,18}$, $d_{14,26}$, $d_{11,18}$ |
| 7 | $d_{1,71}$, $d_{1,161}$, $d_{1,159}$, $d_{1,162}$, $d_{1,68}$, $d_{1,158}$, $d_{1,67}$, $d_{1,75}$, $d_{1,6}$, $d_{1,9}$ |

TABLE S2: Verified functional contacts in cluster 1 arranged starting from the open end of the mouth region towards the hinge region. sc denotes contacts in the side-chain while bb stands for contacts in the backbone.

| Distance | Type | Distance | Type |
|---|---|---|---|
| $d_{20,145}$ | salt bridge | $d_{75,88}$ | hydrophobic |
| $d_{22,141}$ | h-bond sc | $d_{5,60}$ | salt bridge |
| $d_{22,137}$ | salt bridge | $d_{4,64}$ | hydrophobic |
| $d_{21,142}$ | hydrophobic | $d_{4,71}$ | hydrophobic |
| $d_{35,105}$ | hydrophobic | $d_{4,60}$ | hydrophobic |
| $d_{24,106}$ | hydrophobic | $d_{2,64}$ | h-bond sc |
| $d_{22,106}$ | hydrophobic | $d_{7,71}$ | hydrophobic |
| $d_{21,105}$ | h-bond bb | $d_{7,11}$ | h-bond bb |
| $d_{32,105}$ | hydrophobic | $d_{7,12}$ | h-bond bb |
| $d_{30,104}$ | hydrophobic | $d_{8,13}$ | hydrophobic |
| $d_{11,30}$ | hydrophobic | $d_{8,64}$ | salt bridge |
| $d_{20,26}$ | hydrophobic | $d_{4,13}$ | hydrophobic |
| $d_{29,145}$ | hydrophobic | $d_{4,29}$ | hydrophobic |
| $d_{30,145}$ | hydrophobic | | |

*
†

### II. SUPPLEMENTARY FIGURES

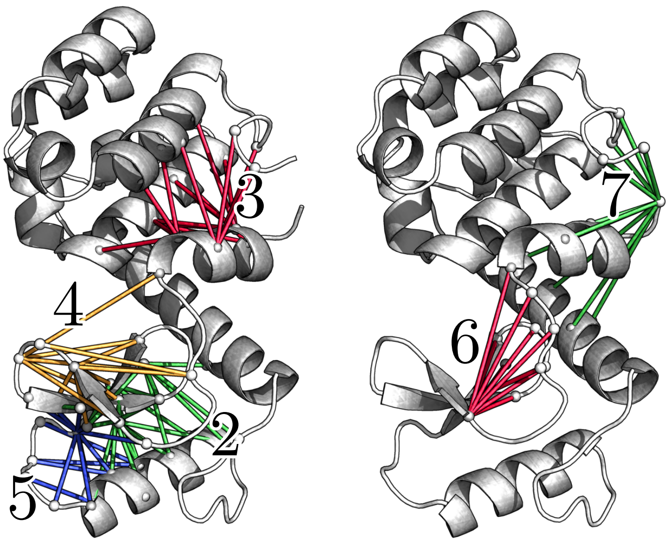


FIG. S1: The next 6 cluster of the correlation matrix after the main open-closed cluster.

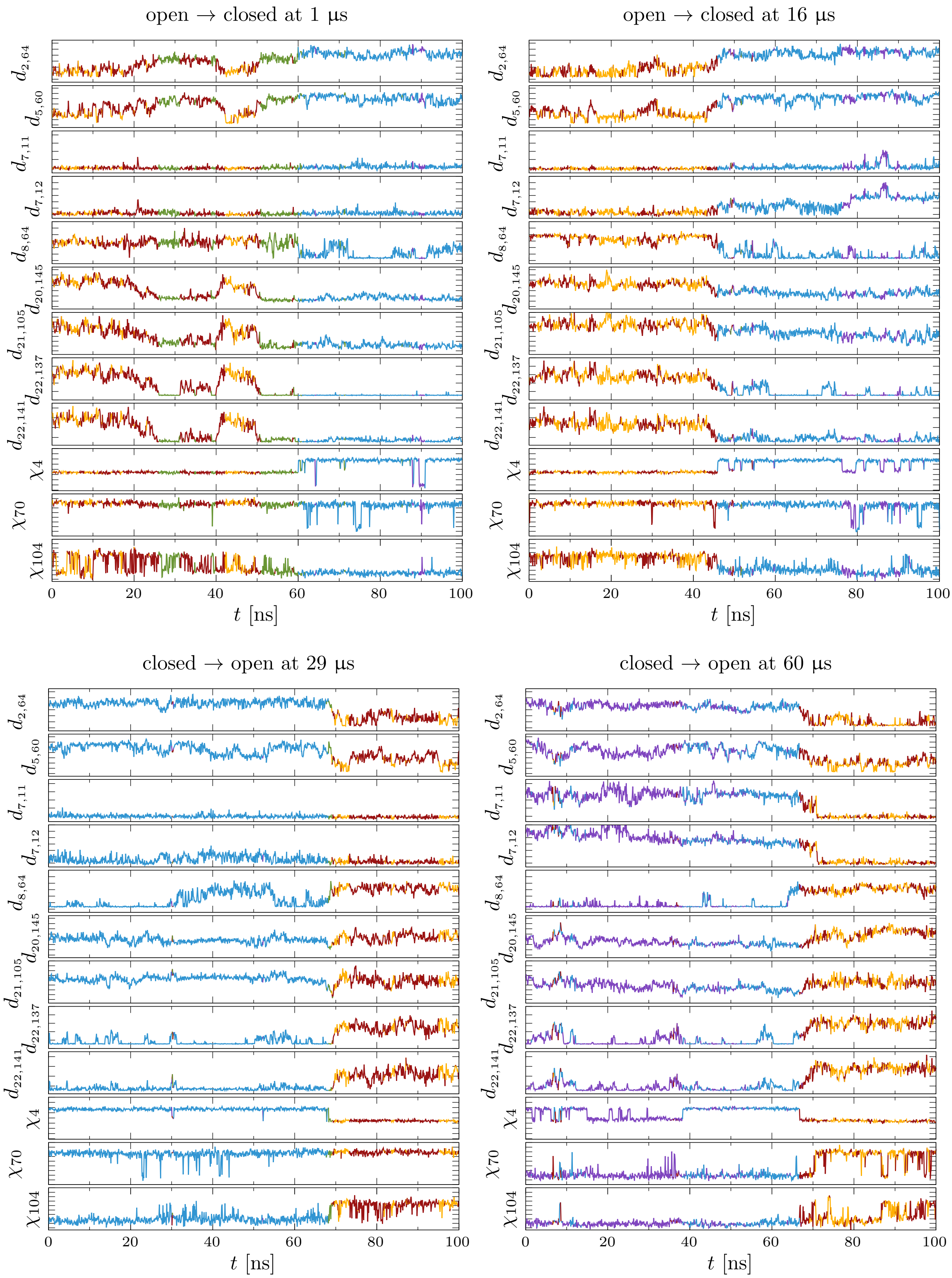


FIG. S2: Time evolution of the selected 12 coordinates of cluster 1 for various open-closed transitions.

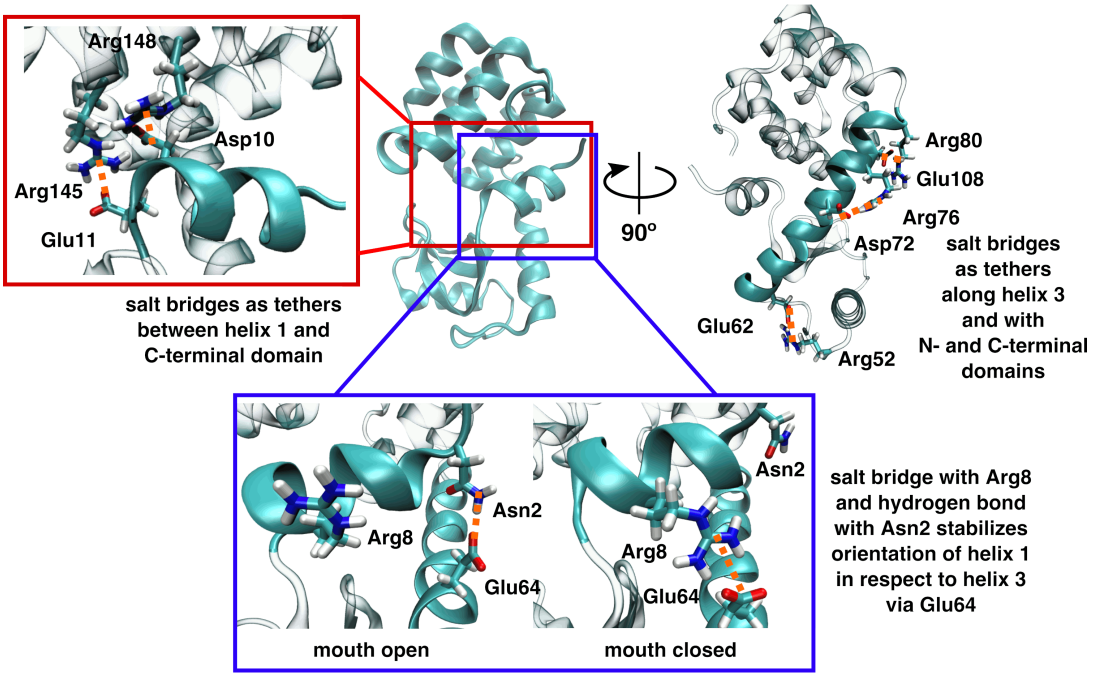


FIG. S3: T4L structural tethers based on salt bridges, showing helix 1/C-terminal domain connection in red inset. A salt bridge complex between Asp10, Glu11, and Arginines 145 and 148 tethers helix 1 to the C domain. While the salt bridge of Arg52 and Glu62 strengthens the connection between helix 3 and the N domain, Arg80 and Glu108 do so between helix 3 and the C domain. The salt bridge between Asp72 and Arg76 strengthens helix 3 above the kink induced by Phe67. Glu64 switches between a hydrogen bond with Asn2 in the mouth open state and a salt bridge with Arg8 in the mouth closed state to stabilize the respective orientations of helix 1 in respect to helix 3.

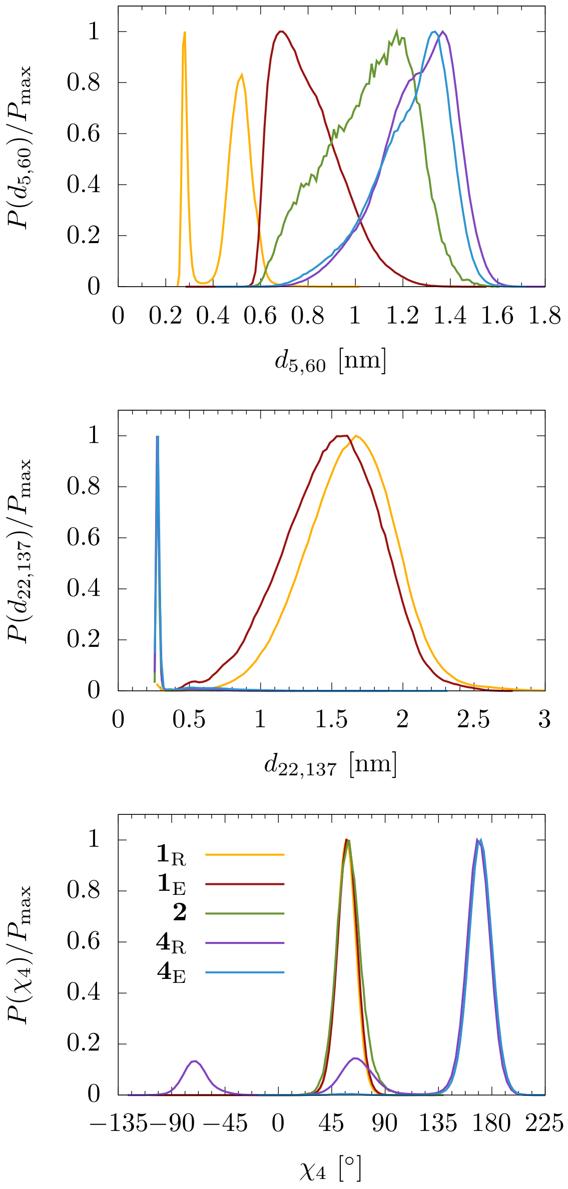


FIG. S4: Marginal probability distribution of the six essential coordinates, corresponding to the five metastable states $\mathbf{1}_\mathrm{R}$, $\mathbf{1}_\mathrm{E}$, $\mathbf{2}$, $\mathbf{4}_\mathrm{R}$ and $\mathbf{4}_\mathrm{E}$.

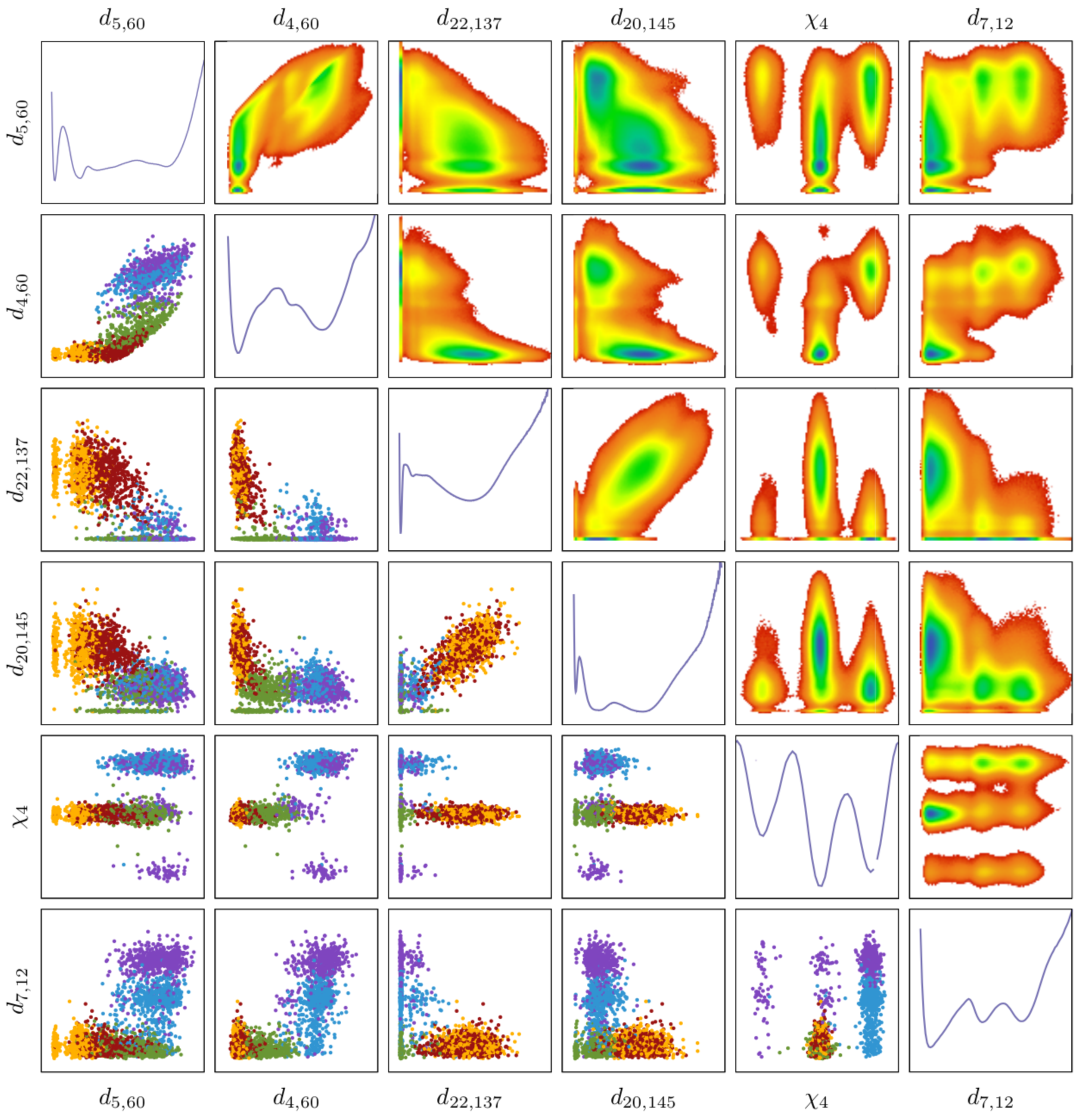


FIG. S5: One- and two-dimensional free energy surfaces along the above defined six coordinates as well as scatter plots of the five metastable states. The color code is: $\mathbf{1}_\mathrm{R}$ (33.5 %) in orange, $\mathbf{1}_\mathrm{E}$ (35.5 %) in red, $\mathbf{2}$ (1.3 %) in green, $\mathbf{4}_\mathrm{E}$ (15.1 %) in blue and $\mathbf{4}_\mathrm{R}$ (14.5 %) in purple.

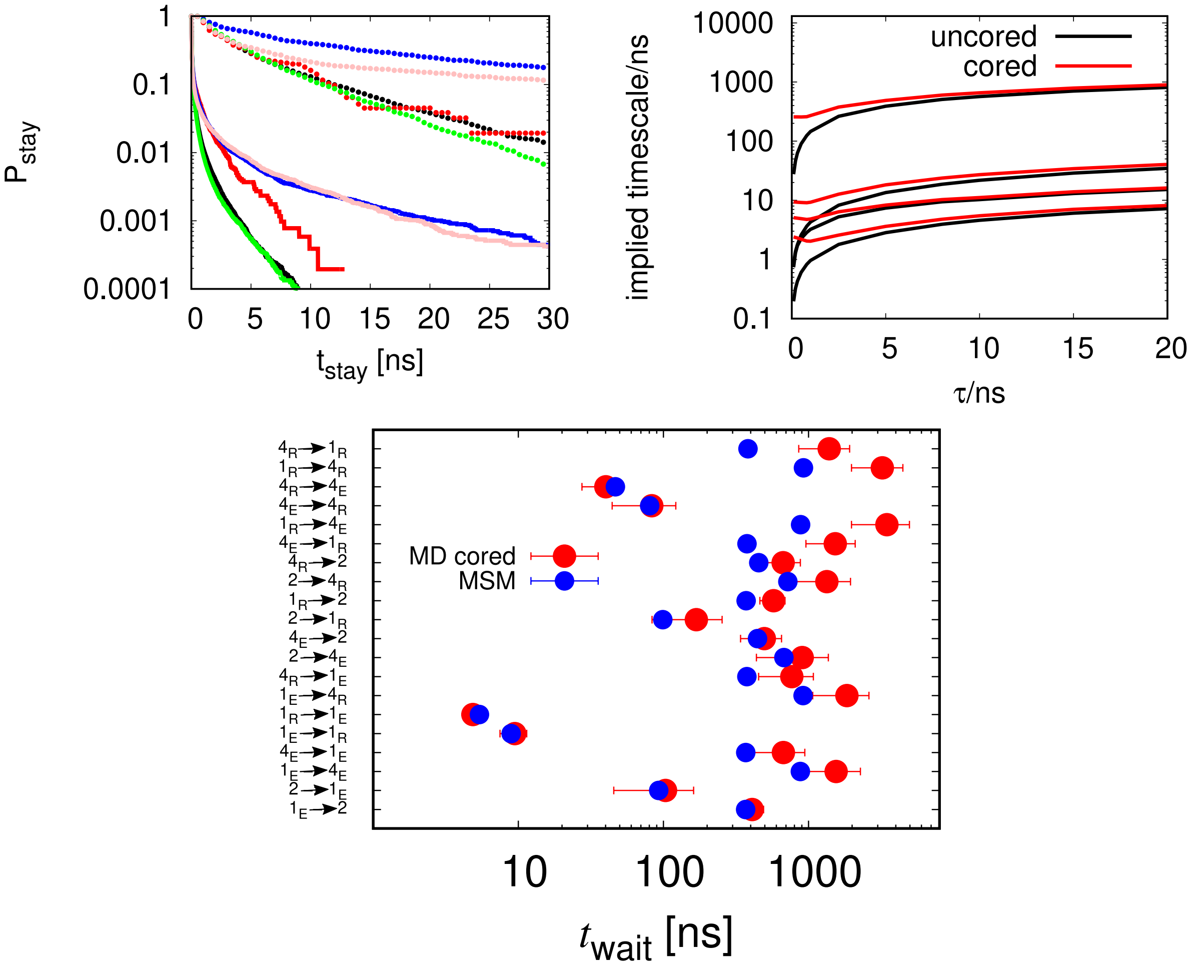


FIG. S6: (Top left) Staying probability of states $\mathbf{1}_{\mathrm{R}}$ (green), $\mathbf{1}_{\mathrm{E}}$ (black), **2** (red), $\mathbf{4}_{\mathrm{E}}$ (blue) and $\mathbf{4}_{\mathrm{R}}$ (pink) are shown. Lines represent the probabilities before coring while dots refer to the cored results with $\tau_{\mathrm{cor}} = 700$ ps. The implied timescales (top right) are shown before (black) and after coring (red). At the bottom, we see the average waiting times of the cored MD data before (red) as well as the predictions of an MSM with lag time $\tau_{\mathrm{lag}} = 700$ ps (blue). The error bars are given as standard deviations of the mean.


[1] V. Traag, L. Waltman, and N. van Eck, "From Louvain to Leiden: guaranteeing well-connected communities", *Sci. Rep.* **9**, 5233 (2019).

[2] G. Diez, D. Nagel, and G. Stock, "Correlation-based feature selection to identify functional dynamics in proteins", arxiv.2204.02770 (2022).
